# The Alzheimer’s Aβ peptide forms biomolecular condensates that trigger amyloid aggregation

**DOI:** 10.1101/2024.01.14.575549

**Authors:** Greta Šneiderienė, Alicia González Díaz, Sourav Das Adhikari, Jiapeng Wei, Thomas Michaels, Tomas Šneideris, Sara Linse, Michele Vendruscolo, Kanchan Garai, Tuomas P. J. Knowles

## Abstract

The onset and development of Alzheimer’s disease (AD) is linked to the accumulation of pathological aggregates formed from the normally monomeric amyloid-β peptide within the central nervous system. These Aβ aggregates are increasingly successfully targeted with clinical therapies, but the fundamental molecular steps that trigger the initial nucleation event leading to the conversion of monomeric Aβ peptide into pathological aggregates remain unknown. Here we show that the Aβ peptide can form biomolecular condensates on lipid bilayers both in molecular assays and in living cells. Our results reveal that these Aβ condensates can significantly accelerate the primary nucleation step in the amyloid conversion cascade that leads to the formation of amyloid aggregates and plaque. We show that Aβ condensates contain phospholipids, are intrinsically heterogenous, and are prone to undergo a liquid-to-solid transition leading to the formation amyloid fibrils. These findings uncover the liquid-liquid phase separation behaviour of the Aβ peptide, and reveal a new molecular step very early in the amyloid-β aggregation cascade that can form the basis for novel therapeutic intervention strategies.

**Significance statement:** The hallmark of Alzheimer’s disease is the abnormal buildup of the normally soluble amyloid β protein aggregates in the central nervous system. While the molecular mechanisms at the late stages of the amyloid β aggregation cascade are well understood, the initial steps remained elusive until now. Our current study demonstrates that amyloid β undergoes liquid-liquid phase separation on lipid surfaces, which triggers primary nucleation and initiates the amyloid β aggregation cascade. This newly identified step in the molecular mechanism of Alzheimer’s disease represents a promising target for the development of alternative innovative therapeutic strategies.

## Introduction

Worldwide over 50 million patients suffer from neurodegenerative disorders, with Alzheimer’s (AD) disease being the most prevalent type (1,2). A central hallmark of AD is the accumulation of extracellular senile plaques and intracellular neurofibrillary tangles in the central nervous system, which ultimately lead to synapse loss, neural cell death and thereby cognitive impairment (1,3). The intrinsically disordered amyloid β1-42 (Aβ42) peptide, which originates from cleavage from the transmembrane amyloid precursor protein (APP), is the most abundant constituent in extracellular amyloid plaques in AD (4). Targeting and removal of these Aβ aggregates is a therapeutic strategy which is increasingly demonstrating successful outcomes in the clinic, especially when administered at an early stage in the disease progression (5). These successes have highlighted the importance of understanding the early events that trigger the conversion of normally monomeric Aβ peptide into pathological aggregates. The later stages of the Aβ aggregation cascade are increasingly well understood, but very little is known about the fundamental mechanisms that underpin the primary nucleation step that initiates this cascade that leads to the conversion of the normal monomeric form of the peptide into pathological aggregates. The monomer may be in a supersaturated state, i.e. kinetically stable above it is solubility limit, as amyloid aggregation is characterised by very significant kinetic barriers which protect the monomeric forms of peptides and proteins from aggregation (6). In particular, there is a very significant nucleation barrier to form the initial aggregates, which, once this barrier is overcome, can grow and multiply more rapidly (7). Recent studies have revealed the key role of biomolecular condensates formed through liquid-liquid phase separation (LLPS) in overcoming the nucleation barrier and triggering pathological aggregation of other proteins involved in neurodegenerative diseases, including tau and ⍺-synuclein. Such condensate systems can in many cases be functional and have important roles as organisational hubs in living cells (8–10). However, the close molecular proximity in such condensates can also promote aberrant interactions leading to aggregation (11–16). However, whether or not the Aβ peptide can form such condensates and whether they are connected to the pathological aggregation pathway has remained unknown. In this study, we used confocal fluorescence and total internal reflection fluorescence (TIRF) microscopy, a set of biophysical characterisation and cell assays to explore the aggregation mechanism of Aβ42 on lipid membranes. We report the discovery of lipid-driven phase separation of Aβ42 *in vitro* and cells. We demonstrate that Aβ42 co-phase separates with phospholipids to form biomolecular condensates. Moreover, these Aβ42 condensates mature, convert into gel-like structures and ultimately grow into amyloid fibrils. These results reveal lipid-driven liquid-liquid phase separation of Aβ42 as a key early step that can trigger AD-linked Aβ42 fibrillation.

## Results

### Aβ42 undergoes liquid-liquid phase separation *in vitro*

An intensive effort has been focused on exploring how different co-factors, such as lipids, which make up to 60% of the dry brain mass (17), affect the rates of distinct Aβ42 fibrillation steps (18–23). Changes in brain lipidome have been linked to the development of AD, giving rise to a great interest in understanding the role of lipid surfaces in Aβ42 aggregation in AD (24). Lipid membranes, depending on their composition, affect Aβ42 fibrillation rates (19,20,23) and there is evidence from kinetic studies that membrane surfaces affect primary aggregation pathways of Aβ42 (19), The molecular mechanisms underpinning this acceleration have, however, largely remained unknown. To address this question, we initially probed Aβ42 aggregate formation on lipid membranes by total internal reflection fluorescence (TIRF) and confocal fluorescence microscopy imaging. Lipids with phosphocholine (PC) headgroups are the most abundant lipid species in mammalian cells (25,26), hence we chose supported lipid bilayers (SLBs) composed of 90% 1,2-dimyristoyl-sn-glycero-3-phosphocholine (DMPC) and 10% cholesterol for our study.

We first explored Aβ42 aggregation on deposited membranes composed of 90% DMPC and 10% cholesterol by TIRF microscopy. At late times, we observed the presence of typical amyloid fibrils observed as the end product of the aggregation process (Figure 1a, b). However, prior to the formation of mature amyloid fibrils, we observed spherical Aβ42 assemblies with diameters in the range of 5–15 μm, and morphologies consistent with biomolecular condensates (Figure 1b). To probe the properties of these assemblies in more detail, we added a solution containing wild-type Aβ42 doped with fluorescently labelled Aβ42 S8C mutant onto a DMPC membrane containing 10% cholesterol and 1% BODIPY™ FL C12-HPC dye in polydimethylsiloxane (PDMS) observation chambers. We found that after 23 hours of incubation, Aβ42 assembled into spherical clusters that colocalise with lipids (Figure 1c). Importantly, Aβ42 does not form such clusters when deposited on a glass slide (Figure S1), indicating that lipids drive the Aβ42 assembly into these species.

**Figure 1.**
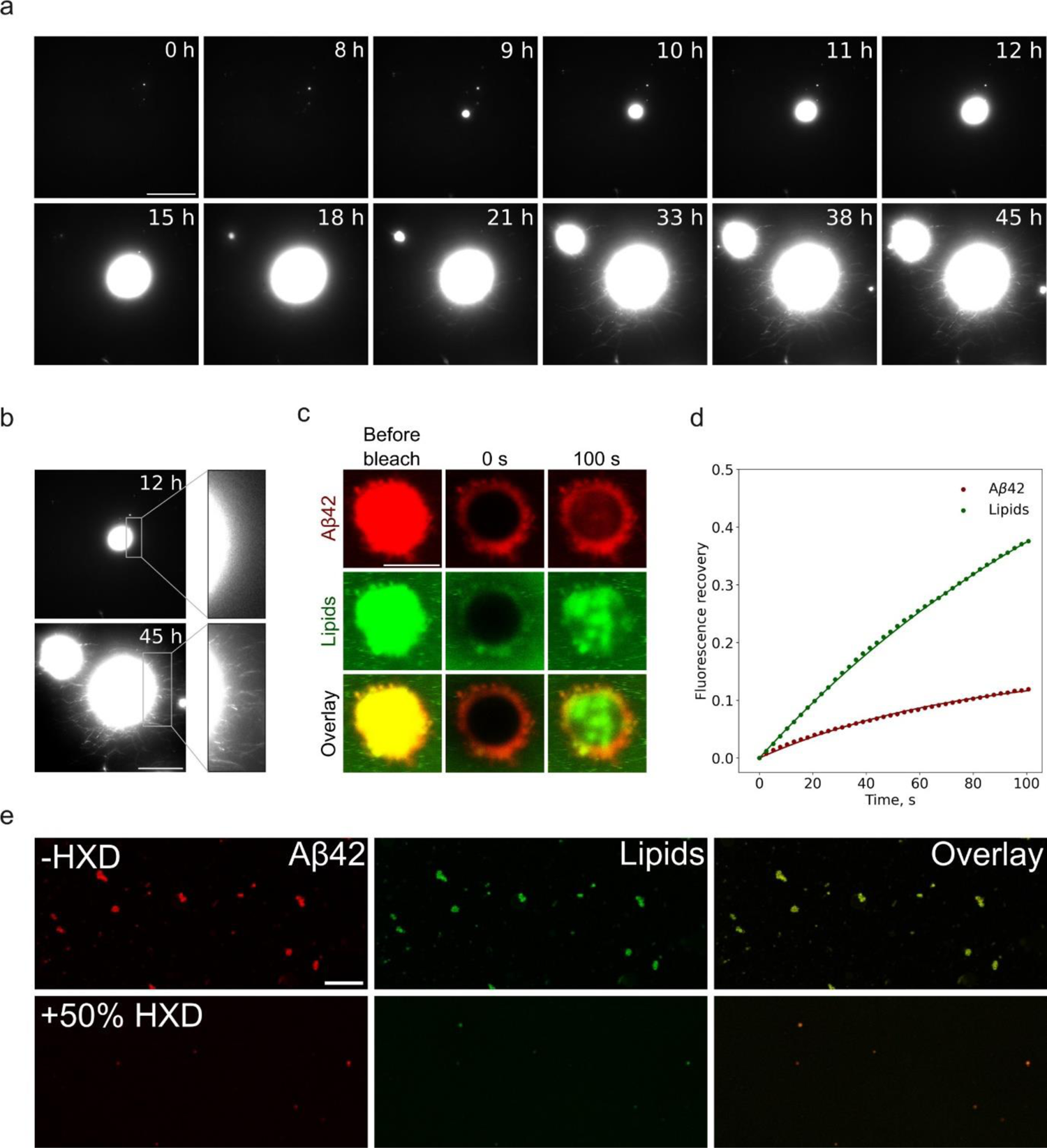
Aβ42 undergoes liquid-liquid phase separation in vitro. a) TIRF images showing the formation of spherical Aβ42 clusters on the DMPC supported lipid bilayer over time. 2 μM Aβ42 and 100 μM lipids were used. Scale bar = 20 μm. b) TIRF microscopy images of ThT-positive Aβ42 condensates after 12 and 45 hours of incubation and insets showing a smooth condensate edge (12 h) and spiked condensate edge containing fibrils (45 h). Scale bar = 20 μm. c) A confocal microscopy image of lipid membrane-colocalised Aβ42 condensates. The time of incubation is 23 h. Scale bar = 5 μm. d) Fluorescence recovery after photobleaching curve of Aβ42 condensates. e) Confocal microscopy images of 1,6-hexanediol (HXD)-treated Aβ42 aggregates. Scale bar = 10 μm.

We next examined the internal dynamics of the clusters by fluorescence recovery after photobleaching (FRAP) experiments. These measurements (Figure 1d) demonstrated rapid recovery in FRAP, which indicates rapid molecular diffusion within the clusters and thus suggests a liquid-like nature of the assemblies as observed for other biomolecular condensates (Figure 1d). Using FRAP, we measured the molecular diffusion coefficients for both the lipid and peptide components. These measurements showed that diffusion coefficient of the protein species in the condensates D = 4.4 ± 1.4 × 10^−14^ m^2^s^−1^ is in the same range as reported in the literature for other biomolecular condensates (27–29). This value is 10^4^ times smaller than the diffusion coefficient of Aβ42 in solution as determined by MDS (D = 1.4 ± 0.1 × 10^−10^ m^2^s^−1^, see methods). The measured diffusion coefficient of the lipids within the condensate D = 2.4 ± 1.3 × 10^−14^ m^2^s^−1^ is of a similar order of magnitude to that found for the peptide in the condensate phase, but lower than the diffusion coefficient of lipids in supported bilayers alone (30,31). Taken together, the diffusion coefficients of protein and lipid species within a condensate are similar and significantly smaller than the diffusion coefficients outside of the condensate environment.

Biomolecular condensates can be stabilised by a rich variety of interactions (32–35), with hydrophobic effect being particularly important. We therefore next sought to elucidate the nature of stabilising interactions of Aβ42 condensates by treating them with a hydrophobic disruptor, 1,6-hexanediol (HXD) (Figure 1e), commonly used to assess biomolecular condensates. Our results show that Aβ42 condensates dissolved in the presence of 50% HXD, supporting the concept that Aβ42 condensates are dense liquid phases assembled as a consequence of the hydrophobic effect.

Overall, these results suggest that Aβ42 undergoes phospholipid membrane-driven liquid-liquid phase separation (LLPS), which is a newly discovered step in the *de novo* Aβ42 fibrillation mechanism. The condensation step occurs at a lipid interface where phospholipids co-assemble with Aβ42 in the condensates.

### Aβ42 undergoes liquid-liquid phase separation in cells

After demonstrating that Aβ42 can undergo biomolecular condensation and that lipids are essential for this process *in vitro*, we set out to study Aβ42 phase separation behaviour on the membrane surface of human neuroblastoma SH-SY5Y cells. The cells were plated and exposed to 2 µM of freshly purified Aβ42 monomer. After 16 h of incubation, the cells were treated with 2.5% w/v HXD, fixed and imaged with a confocal microscope. Both Aβ42 and cell membranes were stained to assess the morphology, number, size and distribution of aggregates on the cellular surface, as shown in Figure 2. First, we noticed that regions without the cells lacked Aβ42 aggregates, suggesting the need for exposed membranes to promote the aggregation of Aβ42 (Figure S2). Secondly, aggregates at early time points after Aβ42 treatment were not irregularly shaped and spiked as typical for fibril clusters but rather appeared as round-shaped structures, typical for liquid biomolecular condensates (Figure 2a, b). By conducting a detailed morphological assessment of observed Aβ42 species, we qualitatively categorised them into two classes (Figure 2b). Class I aggregates were generally small elliptical structures, spread across exposed cell surfaces and mostly co-localised with membranes. By contrast, Class II aggregates were spherical, wrapped around the apoptotic cells and co-localised both with the membrane and DNA-specific dyes.

**Figure 2.**
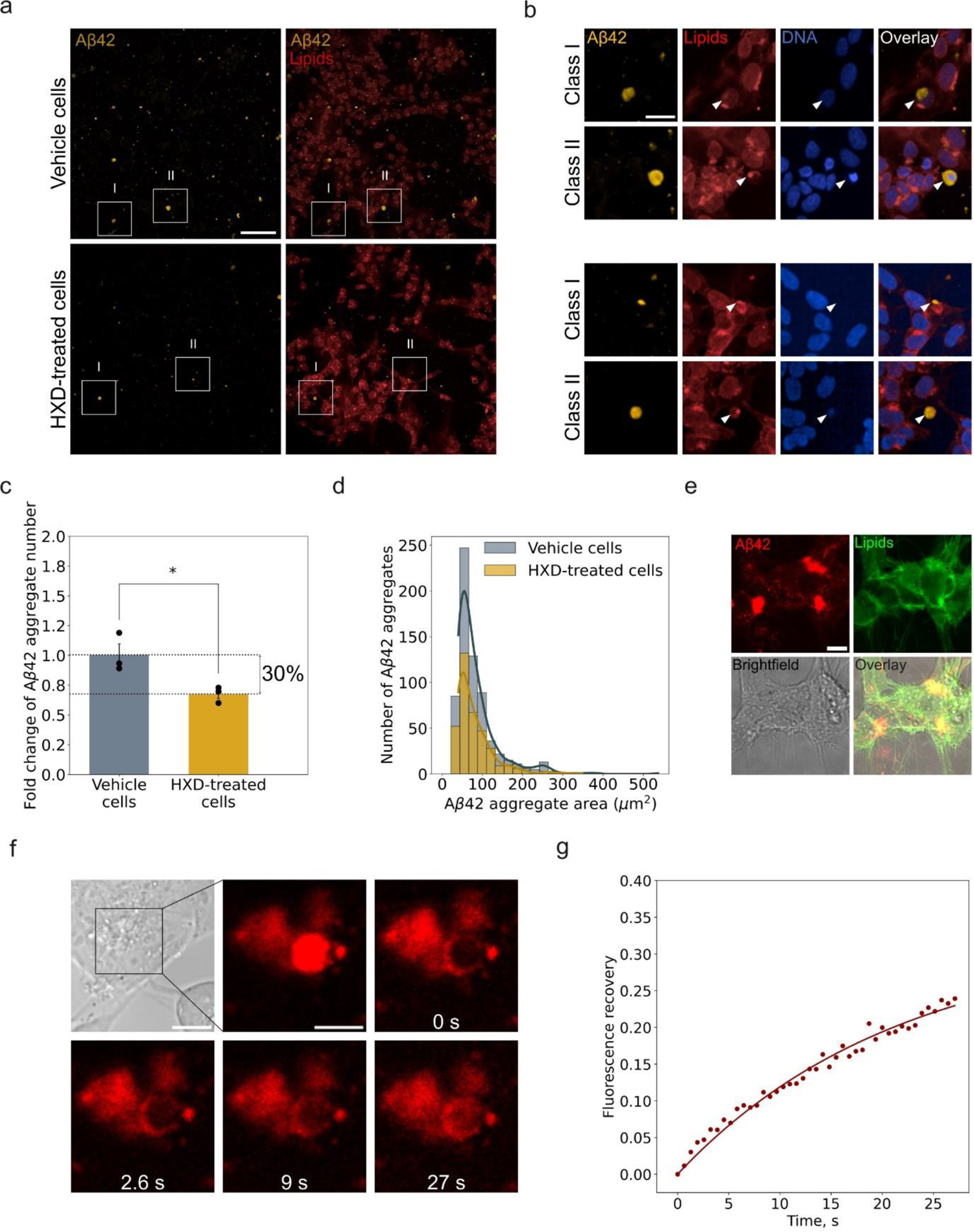
Liquid-liquid phase separation of Aβ42 in human cells. **a)** Representative images of Aβ42 aggregates on human neuroblastoma SH-SY5Y cells treated with vehicle (a, top) or 2.5% w/v 1,6-hexanediol (a, bottom) for 5 minutes. Scale bar = 100 μm. b) Two classes of condensate-like structures are observed in both conditions. Class I aggregates mostly colocalise with the cell membrane and Class II aggregates colocalise with cell membranes and wrap around nucleic acids. Scale bar = 25 μm. The cell membrane was stained with wheat germ-agglutinin, and Aβ42 was stained with W02 anti-Aβ42 antibodies. c) Fold change of the total number of Aβ42 condensate normalised by the total area of SH-SY5Y cells treated with vehicle (VEH) or 2.5% w/v 1,6-hexanediol (HXD). d) Size distribution of Aβ42 aggregates on SH-SY5Y cells (10 000 cells/well density) after treatment with vehicle or 2.5% w/v 1,6-hexanediol. All cell samples were fed with 2 μM Aβ42 monomer and incubated for 16 h before 1,6-hexanediol treatment. Lines show kernel density estimate plots. e) Confocal fluorescence microscopy images of Aβ42 condensates colocalising with cellular membranes. Scale bar = 10 μm. f) Confocal fluorescence microscopy images of Aβ42 condensates recovering after photobleaching. Scale bar = 5 μm. g) Fluorescence recovery after photobleaching curves of the Aβ42 condensates.

Next, we tested the liquid-like properties of Aβ42 aggregates by quantifying the number and sizes of Aβ42 species on 2.5% w/v HXD- and buffer (vehicle, VEH)-treated cells. The total number of Aβ42 objects in both samples was normalized by the total area occupied by cells and plotted in Figure 2c. Cells treated with HXD had a 30% lower Aβ42 aggregate number (Figure 2c) and Aβ42 clusters present in HXD-treated cells were smaller (Figure 2d), indicating that Aβ42 assemblies in SH-SY5Y cells are susceptible to HXD treatment and thereby have liquid-like properties.

The clusters observed in the cell experiments had a similar morphology to the condensates observed *in vitro* and described above. We thus next sought to explore whether the clusters formed on living cells also had a liquid nature as biomolecular condensates. For this purpose, we treated SH-SY5Y cells at 15 000–30 000 cells/well density with 2 µM of freshly purified Aβ42 spiked with fluorescently labelled Aβ42-TAMRA. The cellular membranes were stained with Green CellBrite dye and imaged after 16–24 hours after treatment (Figure 2e). The Aβ42 assemblies on SH-SY5Y cells colocalised with the lipids, in agreement with our *in vitro* observations (Figure 1c). The fluorescence signal of Aβ42 condensates recovered after photobleaching, indicating that Aβ42 assemblies are indeed liquid-like (Figure 2f, g). Overall, we provide evidence that Aβ42 associates with cell membranes and forms liquid-like, HXD-sensitive condensates containing both protein and lipid phases in SH-SY5Y cells.

### Aβ42 condensates undergo a liquid-to-solid transition *in vitro*

We next turned to probe whether the Aβ42 condensates were connected to the disease-associated amyloid aggregation pathway of the peptide. To explore the properties of the condensates throughout their lifetime, we tested the dependence of Aβ42 condensate susceptibility to HXD and KCl treatment on condensate age (Figure 3a, b). Early stage (3.5 h) Aβ42 condensates were soluble in 100% HXD, whilst mature (22 h) condensates remained intact upon treatment with 100% HXD (Figure 3a top and bottom, respectively). Notably, both early-stage and mature condensates are insoluble in high salt solution (0.5 M KCl). These results suggest that early stage Aβ42 condensates are a consequence of the hydrophobic effect rather than electrostatic interactions and undergo a liquid-to-solid transition upon maturation.

**Figure 3.**
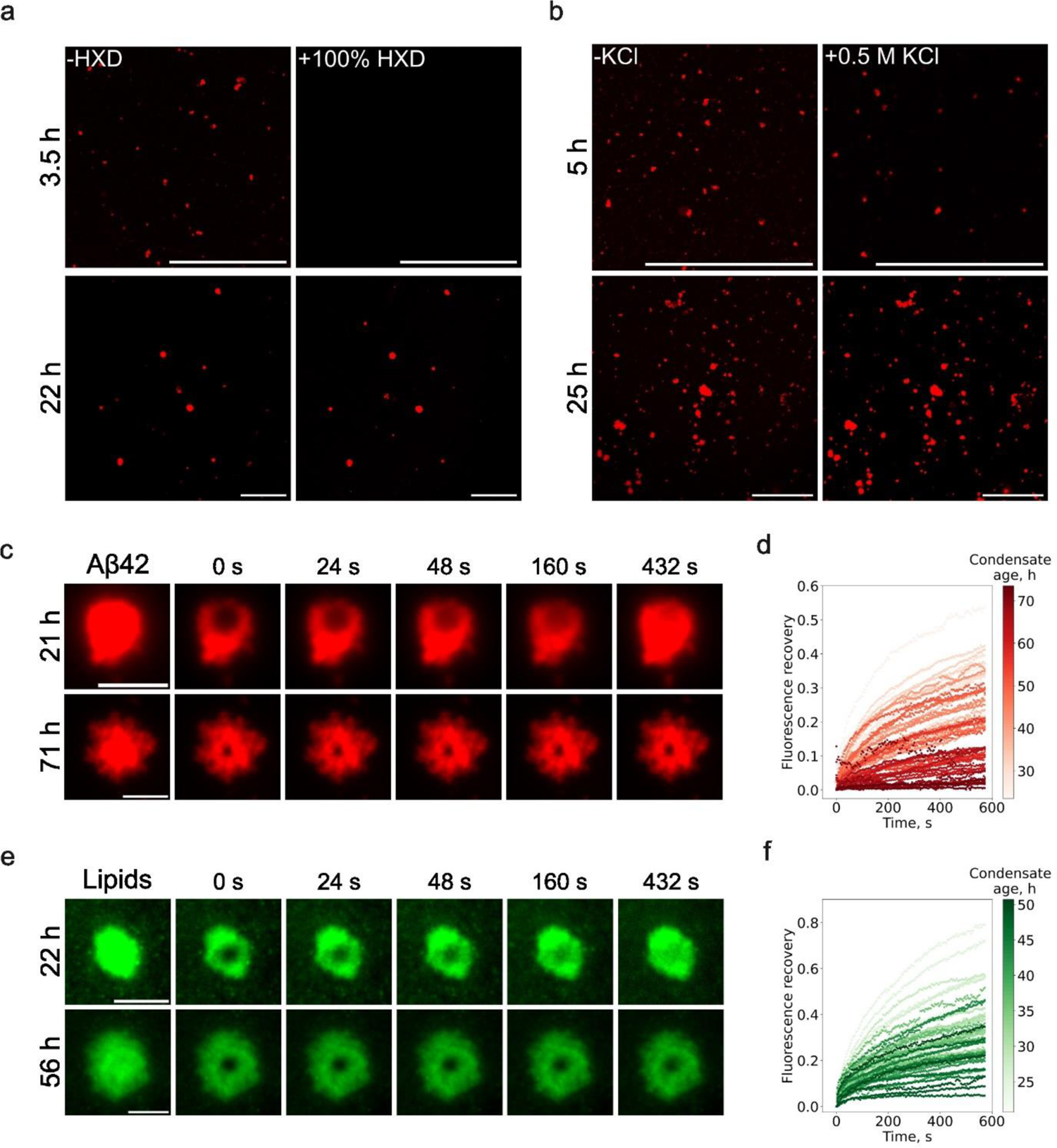
Aβ42 condensates undergo a liquid-to-solid transition on DMPC membranes *in vitro*. Confocal microscopy images of Aβ42 condensates treated with 100% (6 M) 1,6-hexanediol (a) and 0.5 M KCl (b). Aβ42 condensates were incubated for 3.5–22 h before 1,6-hexanediol and 5–25 h before KCl treatment. Scale bar = 50 μm. c) TIRF microscopy images of ageing Aβ42 aggregates during the FRAP experiment (Aβ42 imaging channel). Scale bar = 5 μm. d) FRAP recovery dependence on Aβ42 condensate maturation stage (Aβ42 imaging channel). e) TIRF microscopy images of ageing Aβ42 aggregates during the FRAP experiment (lipid imaging channel). Scale bar = 5 μm. f) FRAP recovery dependence on Aβ42 condensate maturation stage (lipid imaging channel).

We also noted that lipids have different responses to HXD treatment depending on whether they are inside or outside of the condensate environment. Lipids, that colocalise with 5% HXD-resistant Aβ42, remain in the condensates after 5% HXD treatment, while free lipids and lipids that are inside of 5% HXD-sensitive condensates are solubilised upon 5% HXD treatment (Figure S3, right, white arrows). This suggests that lipids and Aβ42 mature together and age by forming a uniform HXD-resistant phase.

To explore the internal mobility properties of ageing Aβ42 condensates, we measured FRAP recoveries on condensates at their different maturation stages (Figure 3c, d, e, f). The rate of fluorescence recovery in the protein fraction correlates with the condensate age (Figure 3d). Mature Aβ42 condensates demonstrate a lower fluorescence recovery rate than fresh ones (Figure 3d), suggesting that Aβ42 mobility inside a condensate becomes more restricted over time. This is also true for the lipid fraction in the condensates, although the correlation between FRAP recovery rate and condensate age is lower (Figure 3f).

Overall, our findings indicate that Aβ42 condensates are metastable assemblies. Aβ42 initially forms liquid drops during lipid membrane-driven phase separation, these assemblies then grow by association of free Aβ42 monomer from the solution and eventually turn into gel-like and solid particles over time.

### Aβ42 condensates are heterogenous *in vitro* and cells

As we discovered that Aβ42 condensates mature over time and undergo a liquid-to-solid transition, we then set out to explore the internal structure of individual condensates. For this purpose, we bleached two different zones in the same condensate: the centre (Figure 4a, top) and the edge (Figure 4a, middle) and then recorded the fluorescence recovery curves. Fluorescence recovered quickly in the central zone of the condensate, while the bleached edge of the condensate remained dark (Figure 4a, b). This suggests that Aβ42 condensates are internally heterogenous and co-exist in liquid and solid states with a liquid centre and solid edge. Hence different parts of the same Aβ42 droplet have different relaxation properties, suggesting that condensate maturation rates vary across the condensate. Our findings further indicate that Aβ42 condensate maturation starts at the interface of the condensate. At late stages of maturation, fully fibrillar aggregates are visible on the outer interface of the condensate (Figure 1b, S6). This type of interface activity is in agreement with observations on different condensate systems associated with protein aggregation disorders, including FUS and hnRNPAI protein systems (13,36), and suggests that an aggregation-promoting interface could be a general feature of the condensed phases of proteins.

**Figure 4.**
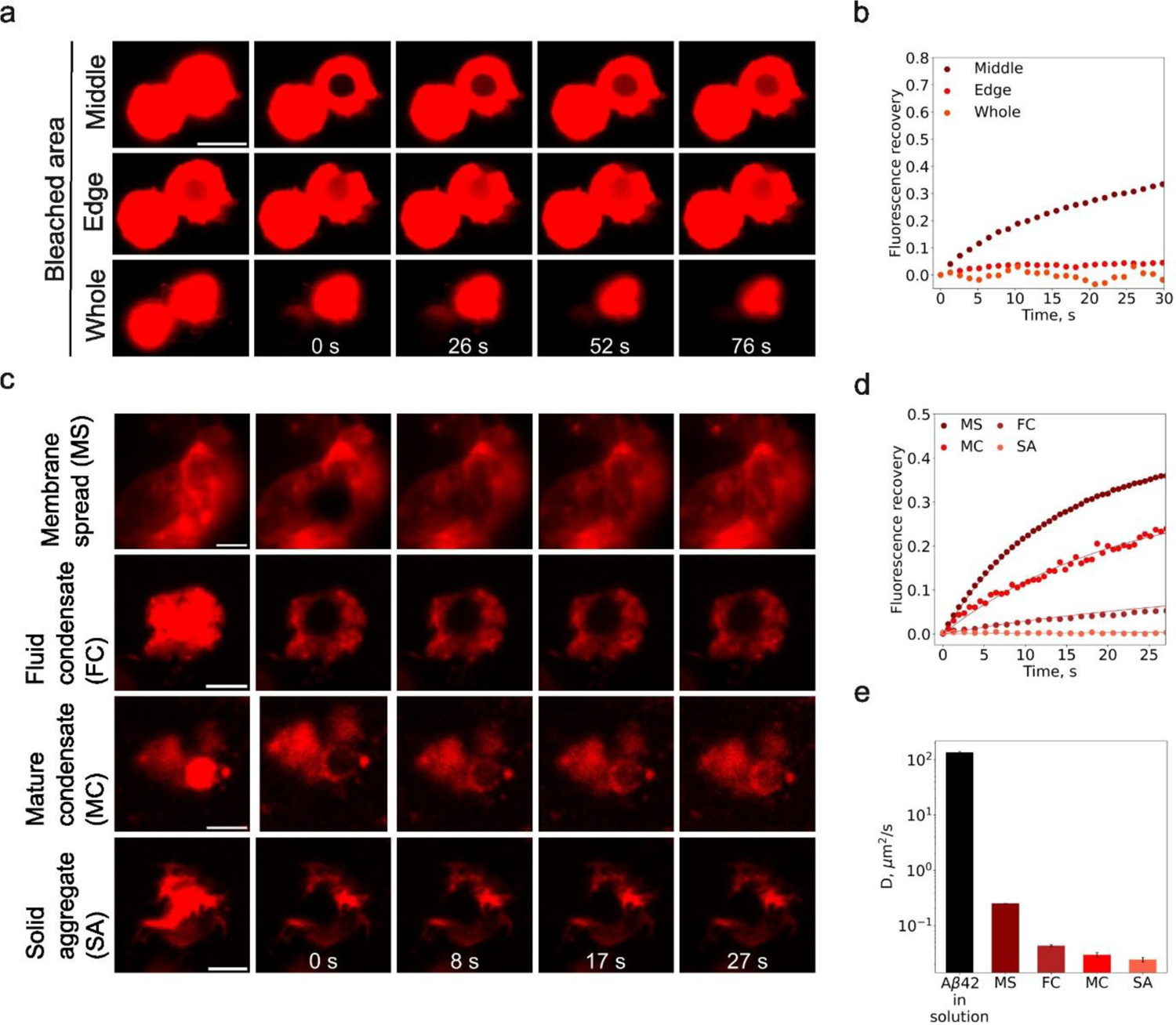
Spatial heterogeneity of Aβ42 condensates *in vitro* and cells. a) Confocal microscopy images of an Aβ42 condensate before and after FRAP recovery after bleaching different zones of the condensate. 2 μM Aβ42 was incubated on DMPC + 10% cholesterol supported lipid bilayer for 25 h before imaging. Scale bar = 5 μm. b) FRAP recovery curves in different zones of the condensate in a). c) Examples of different types of Aβ42 assemblies on SH-SY5Y cells: Aβ42 spread on the membrane (top), fluid Aβ42 condensate (2^nd^ row from the top), mature Aβ42 condensate (3^rd^ row from the top), solid Aβ42 aggregate (bottom row). Scale bar = 5 μm. d) FRAP recovery curves of different Aβ42 species. e) Diffusion coefficients of distinct condensate species, determined by FRAP (MS, FC, MC, SA) and MDS (Aβ42 in solution).

Notably, we did not observe full fluorescence recovery in FRAP experiments when the entire condensate was bleached (Figure 4a, bottom). This finding indicates that material exchange between condensate and the dilute phase surrounding it is decreased due to the solid outer shell.

To further explore the internal structural features of Aβ42 condensates formed in cells, we performed FRAP experiments on individual condensates and qualitatively categorized them into four classes based on fluorescence recovery rates after FRAP, diffusion coefficients and morphology (Figure 4c, d, e). Aβ42 assemblies can occur as a protein spread over an extended area in the cell, with a high fluorescence recovery rate and a diffusion coefficient comparable to diffusion coefficients of protein multimers diffusing laterally in the lipid membrane (Figure 4c, top) (37). Other Aβ42 assemblies include smaller, irregularly shaped structures with liquid-like (fluid condensate, FC) and gel-like (mature condensate, MC) central zone with diffusion coefficients in the 0.03–0.04 μm^2^/s range. Finally, a subpopulation of the condensates appeared as spiky aggregates which had no fluorescence recovery after photobleaching (Figure 4c, bottom, Figure 4d, e). All of these assemblies were present at late incubation time points, which indicates that Aβ42 phase separation and subsequent aggregation are dependent on the local microenvironment in the cells.

Our findings thus suggest that Aβ42 condensates display a range of morphologies and physical properties *in vitro* and cells and can show liquid-solid coexistence during the maturation process. Furthermore, we provide evidence that Aβ42 condensate ageing starts at the interface of dense liquid condensed phase rather than uniformly across a whole condensate.

### Aβ42 condensates mature into aggregates in cells

We discovered that Aβ42 undergoes lipid-driven phase separation in SH-SY5Y cells and evidence from *in vitro* assays suggested that Aβ42 condensates are transient species which convert into solid structures over time. In light of these findings, we tested whether Aβ42 condensate maturation into solid aggregates also occurs in the cellular environment.

SH-SY5Y cells were plated at very low density (1/8 of confluency), treated with 2 µM Aβ42 monomer and incubated for 0, 6, 16, 24 and 48 h at 37°C. At each time point, cells were treated with HXD, stained with membrane- and Aβ42-specific antibodies and fixed. At early time points, 0 and 6 h, we did not observe any Aβ42 aggregates in any field of view (Figure 5a). A low intensity Aβ42 signal was present at the edge of cells after 6 h of incubation, indicating Aβ42 monomer adsorption on the cellular membranes (Figure 5c). Notably, after 16 h of incubation, Aβ42 aggregates of higher intensity, wrapping around or extending on membranes, were observed (Figure 2b, Class II aggregates). Moreover, the number of those aggregates decreased upon HXD treatment, as previously reported (Figure 2c, d and 5b). After longer incubation times, irregular Aβ42 assemblies with protruding fibrils following membrane surfaces were observed (Figure 5a, c). Furthermore, aged aggregates were less sensitive to HXD treatment, in agreement with the *in vitro* data discussed above and indicating the liquid-to-solid transition of Aβ42 condensates (Figure 5b).

**Figure 5.**
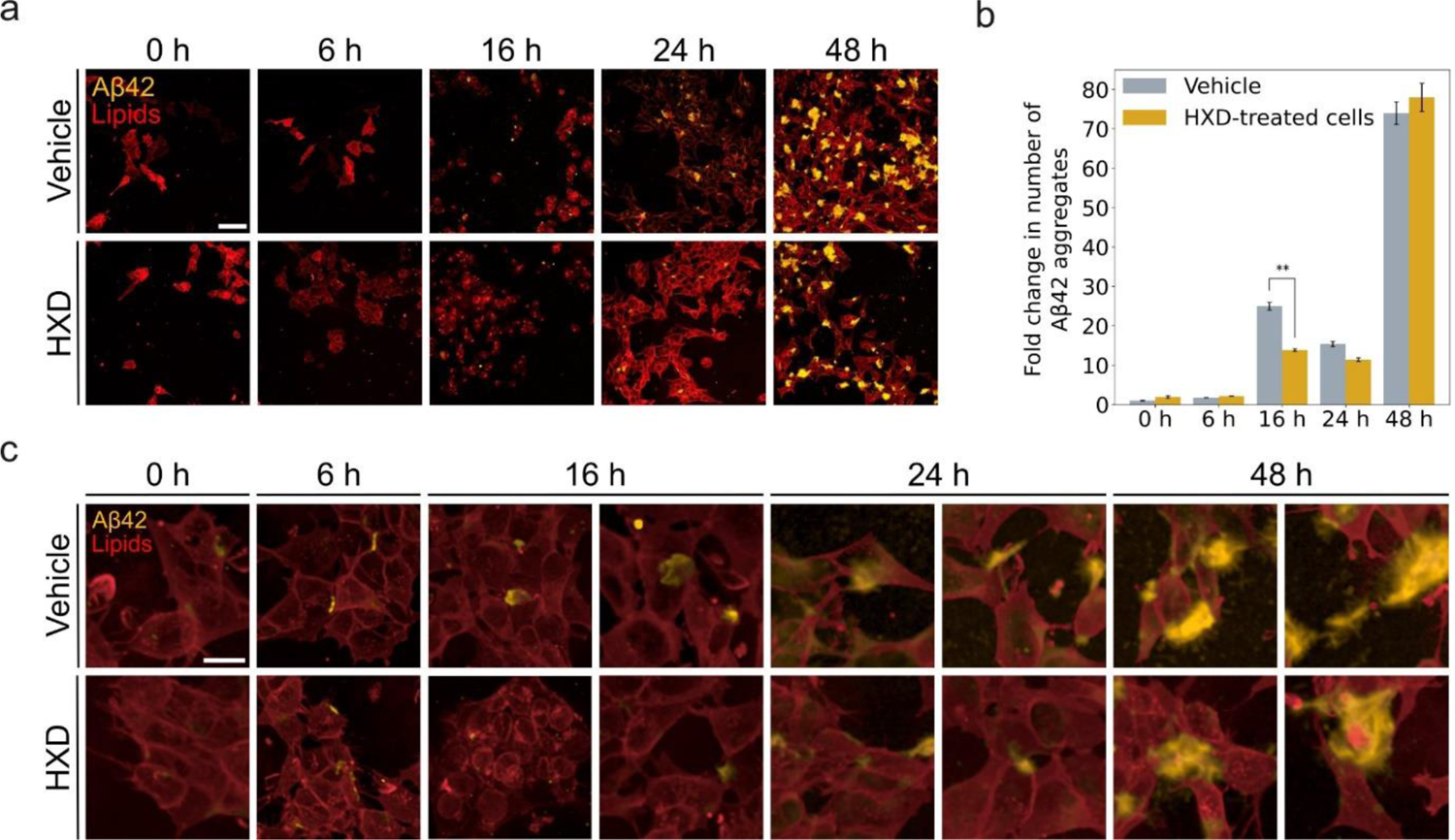
Maturation of Aβ42 condensates in human cells. Progression of Aβ42 aggregates over time upon exogenous addition of 2 μM of monomer to human SH-SY5Y cells. a) Representative confocal microscopy images of cells after 0, 6, 16, 24 and 48 hours of incubation with the protein, treated with vehicle or HXD for 3 minutes before fixation and staining. Scale bar = 100 μm. b) Fold change increase over time of the number of Aβ42 aggregates measured after treatment with vehicle or HXD. Cell density 5 000 cells/well. c) Images illustrating the maturation of Aβ42 aggregates upon interaction with lipid membranes. After 16 hours of incubation, round-shaped aggregates are observed. Spherical aggregates with irregular spikes are observed after 24 and 48 hours, and they become more resistant to hexanediol treatment. Scale bar = 25 μm. HXD – 1,6-hexanediol.

More Aβ42 aggregates formed upon Aβ42 treatment at low cell densities (5 000–10 000 cells/well) compared to high cell densities (20 000–40 000 cells/well) (Figure S4a, b). This effect is attributed to a high Aβ42:membrane ratio. At low cell density (high Aβ42:membrane ratio), a large amount of Aβ42 is adsorbed on a small exposed area of cells, which increases the rate of condensate formation and subsequent maturation into fibrillar structures. By contrast, at high cell densities, Aβ42 is spread over a large area of the membrane surface, which results in larger spacing between Aβ42 monomers and thereby lower aggregation rates. Furthermore, at low cell densities, Aβ42 forms a smaller number of condensates due to the more limited membrane surface area, but these condensates grow to reach a larger size due to the higher level of monomer available to be incorporated into each condensate nucleus (Figure S4c, left). By contrast, a large surface area at high cell densities allows Aβ42 to nucleate at multiple sites, but these nuclei then have a more limited ability to grow due to Aβ42 monomer depletion from the solution and thus remain small (Figure S4c, right). Accordingly, at high cell densities, we mostly detected thousands of small Aβ42 species, with an area <50 µm^2^ (Figure S4c, right). By contrast, at low cell densities, Aβ42 aggregates were much larger, with areas ranging from 50 to > 400 µm^2^ (Figure S4c, left).

Taken together, these data show that Aβ42 condensation dynamics depend on the local microenvironment and the Aβ42:cell ratio. This ratio, which directly translates into the Aβ42:membrane surface area ratio, also impacts the Aβ42 condensate size and number. Moreover, Aβ42 condensates formed in SH-SY5Y cells, mature over time, as indicated by the reduced condensate susceptibility to HXD treatment.

### Thermodynamics of Aβ42 LLPS and aggregation

After the discovery of Aβ42 LLPS *in vitro* and cells, we next sought to gain an understanding of the driving forces behind Aβ42 LLPS and aggregation. To this effect, we quantified the free energy changes governing these processes. To achieve this objective, we measured the concentration of the monomeric peptide in equilibrium with the liquid and solid dense phases by performing FCS measurements with a TIRF setup equipped with an FCS module (38). The fluorescence autocorrelation data were converted to the absolute concentration of free monomeric Aβ42 in solution using the fluorescence autocorrelation of monomeric Aβ42 data as a reference (Figure 6d, S5). Our observations revealed a significant drop in the concentration of monomeric Aβ42 from 2 μM to ca. 300 nM during the initial 37 hours under LLPS-promoting conditions (Figure 6d). Subsequently, the concentration of the free monomer stabilized and remained at approximately the same level. This observation suggests that Aβ42 condensates nucleate and grow during the first 37 hours, while the concentration of monomeric Aβ42 is higher than the saturation concentration of Aβ42 (c_sat_) with respect to phase separation. When the concentration of Aβ42 monomer approaches c_sat_, the condensate growth rate becomes very low and the system reaches equilibrium, with a concentration of 314 nM of free monomeric Aβ42. This value corresponds to a standard Gibbs free energy gain ΔG°_liquid_ = −36.6 kJ/mol for monomer assembly in the condensate.

**Figure 6.**
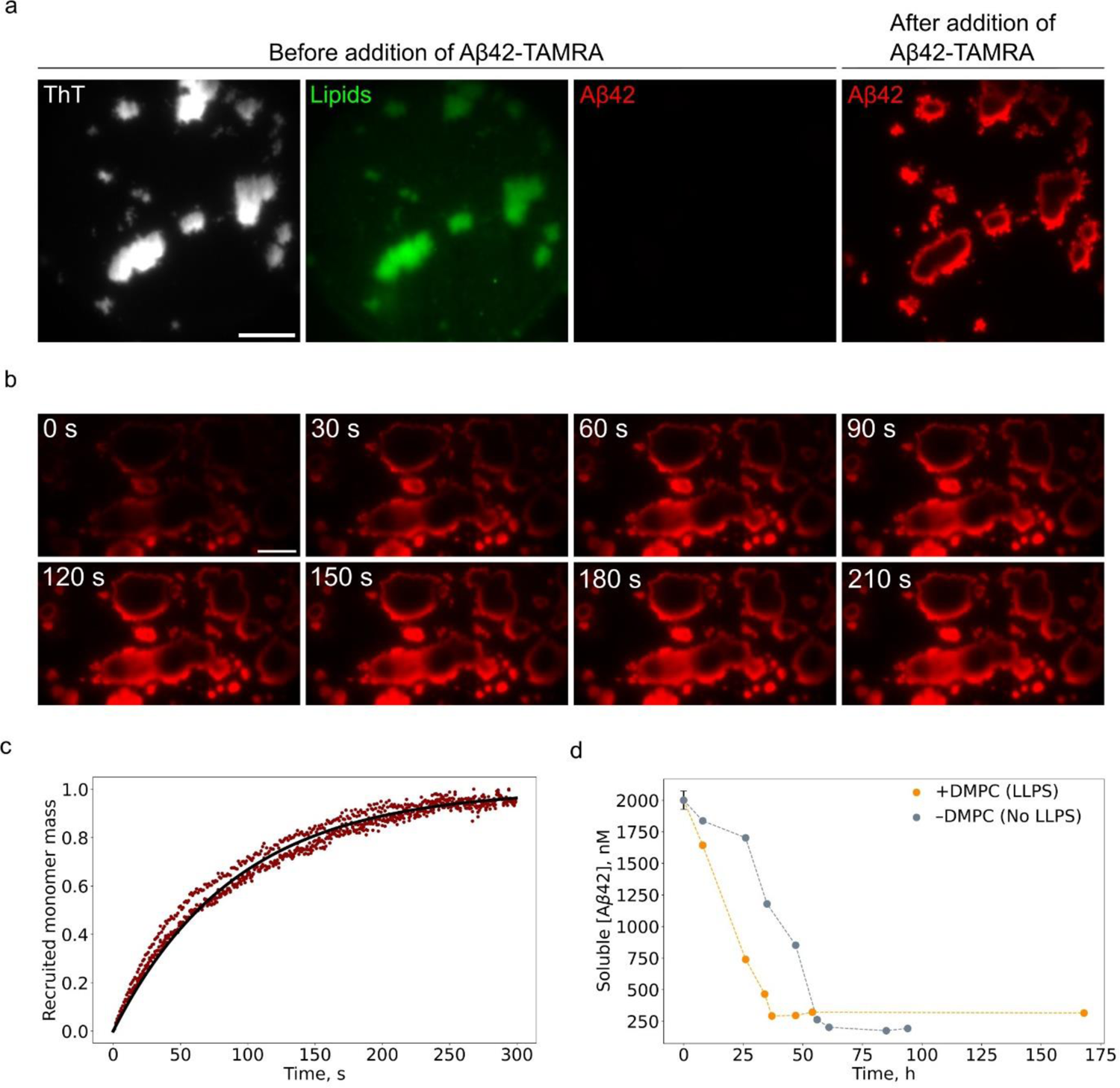
Kinetics of Aβ42 condensate growth. **a)** TIRF microscopy images of Aβ42 condensates before and after the addition of monomeric Aβ42-TAMRA. Monomeric unlabelled Aβ42 was incubated for 48 hours on DMPC supported lipid bilayer before the addition of 100 nM Aβ42-TAMRA. Scale bar = 10 μm. b) TIRF image series demonstrating Aβ42-TAMRA assembly into condensates. Scale bar = 10 μm. c) Kinetics of Aβ42 association into condensates. The black line represents a diffusion-limited Aβ42 association model. N = 3 curves are shown; each curve represents an independent repeat. Monomeric Aβ42 depletion kinetics: d) Change of Aβ42 monomer concentration over time under LLPS and non-LLPS conditions. Dashed lines are guides to the eye. [Aβ42 WT] at t = 0 is 1.9 μM, [Aβ42-ATTO488] = 0.1 μM.

Similarly, under non-LLPS conditions (without a lipid membrane) the concentration of monomeric Aβ42 dropped from 2 μM to 201 nM during the first 61 hours (Figure 6d). Subsequently, the concentration of Aβ42 in solution remained constant at ca. 174 nM, which aligns well with the solubility limit of Aβ42 (30 nM) with respect to amyloid formation determined previously (39). This value corresponds to a standard free energy ΔG°_solid_ = −38.1 kJ/mol Gibbs free energy gain for the transition between monomeric and fibrillar states. The Gibbs free energy difference between liquid and fibrillar states is therefore 1.5 kJ/mol. Notably, our findings highlight that liquid condensed Aβ42 is in a higher Gibbs free energy state than fibrillar Aβ42, indicating system’s progression towards lower energy states during Aβ42 condensation and fibrillation. This aligns with the concept that the amyloid state represents the most stable configuration for a protein above the solubility limit (6,40). As such, condensation can serve as an intermediary step in the Aβ42 fibrillation cascade and is an example of Ostwald’s rule of stages (41,42).

### Kinetics of Aβ42 condensate growth

After having determined the thermodynamic stabilities of the liquid and solid phases, we next focused on the kinetics of the transitions to and from these phases. We initially probed the dynamics of monomer assembly into condensates. For this purpose, we formed ThT-positive, lipid-colocalising Aβ42 condensates (Figure 6a), introduced 100 nM Aβ42-TAMRA and followed association of the labelled monomer to the pre-formed condensates by measuring fluorescence intensity over time (Figure 6b, c). Upon introduction of Aβ42-TAMRA, we observed a gradual partitioning of Aβ42-TAMRA from solution to the edges of the preformed condensates, followed by monomer diffusion towards the centre of the condensates (Figure 6b). The fluorescence intensity time trace, which directly translates into the mass of associated monomer, had a sigmoidal shape (Figure 6c). Condensate growth gradually slowed down and saturated after ca. 250 s, suggesting that Aβ42 condensate growth is limited when the available monomer is depleted. We thus fitted the condensate growth kinetics data with first order reaction rate equation:

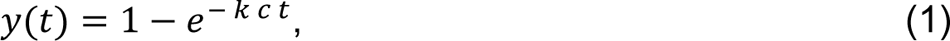

where y is the associated Aβ42 monomer mass, t is time and k is the rate constant for Aβ42 monomer association as a fitted parameter. From the fit we obtained an Aβ42 association into condensates rate constant k = 2.7·10^4^ M^-1^s^-1^. This value is of a similar order of magnitude as the Smoluchowski diffusion-limited protein association rate constant (r = 4πDR = 10^4^–10^6^ M^-1^s^-1^) (43,44). As such, the thermodynamic barriers for Aβ42 monomer association into condensates are small, in agreement with the behaviour of liquid-liquid phase separated systems. This barrier can be estimated from:

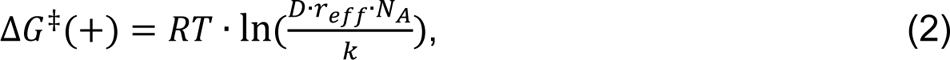

where D = 1.4 ± 0.1 × 10^−10^ m^2^s^−1^ is the diffusion coefficient of free Aβ42 in solution, as determined by MDS; r_eff_ = 0.5 nm corresponds to the effective reaction volume radius of Aβ42 monomers, as estimated in (45); C_dilute_ = 414 nM for the free monomeric Aβ42 in the LLPS state. For the liquid to solid transition, we take the value of 40 mg/ml determined for other phase separating protein systems (47) corresponding to C_dense_ = 8.9 mM. N_A_ is the Avogadro’s number, and k is the monomer association rate constant. Given these parameters, the Gibbs free energy barrier for monomer association is ΔG^‡^(+) = 18 kJ/mol. Notably, the monomer association with condensates Gibbs free energy barrier corresponds to the monomer association process with aged, possibly gel-like condensates (>35 h). Hence, the free energy barrier for monomer association with newly formed liquid condensates may be even lower. Together with the value determined for the overall free energy change ΔG = RT ln(C_dense_/C_dilute_) = RT ln(8.9 mM/414 nM) = 24 kJ/mol, we can evaluate free energy barrier for monomer dissociation from the condensates as ΔG^‡^(-) = 24 + 18 = 42 kJ/mol, which is significantly larger than the Gibbs free energy barrier for monomer association, reflecting the stability of the condensates.

### Biomolecular condensates of Aβ42 of accelerate the primary nucleation step in the aggregation cascade

The experimental observations show that under conditions where LLPS was observed, aggregates appeared within the condensates earlier than under non-LLPS conditions in absence of a lipid membrane (Figure S6). This hinted at differences in the Gibbs free energy barriers for transitioning from free monomeric to fibrillar and condensate states. We can convert the characteristic timescales for the transitions to an energy barrier (7):

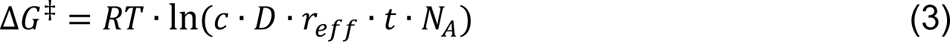

For all phase transitions considered in this study, the parameters are D = 1.4 ± 0.1 × 10^−10^ m^2^s^−1^ corresponding to the diffusion coefficient of free Aβ42 in solution, as determined by MDS; r_eff_ = 0.5 nm is the effective reaction volume radius of Aβ42 monomer, as estimated in (45) (see Materials and Methods for more details). The characteristic timescale for the monomeric to phase separated transition t = 3 h can be estimated from confocal microscopy experiments (Figure S7) and c = 2 μM is the concentration of free monomeric Aβ42 before the transition; N_A_ is Avogadro’s number. For the direct solution to amyloid fibril transition, the timescale can be estimated from Figure 6d as t = 61 h. For the liquid to solid transition, the characteristic timescale can be determined by TIRF microscopy imaging (Figure 1a) as t = 21 h and c = 314 nM is the concentration of free monomeric Aβ42 in dilute phase. Using the above mentioned parameters in Eq. (3) we obtain ΔG^‡^_fib_ = 40.9 kJ/mol for transition between monomeric and fibrillar states, ΔG^‡^_LLPS_ = 33.5 kJ/mol for monomeric to liquid transition and ΔG^‡^_LTS_ = 33.7 kJ/mol for liquid to solid transition (Figure 7a). Importantly, the free energy barrier for Aβ42 liquid-liquid phase separation (33.5 kJ/mol) is lower than that for direct Aβ42 fibrillation (40.9 kJ/mol). This finding suggests that the lipid membrane reduces the energy barrier during the initial phase of Aβ42 assembly and directs the protein to the liquid-liquid phase separation pathway.

**Figure 7.**
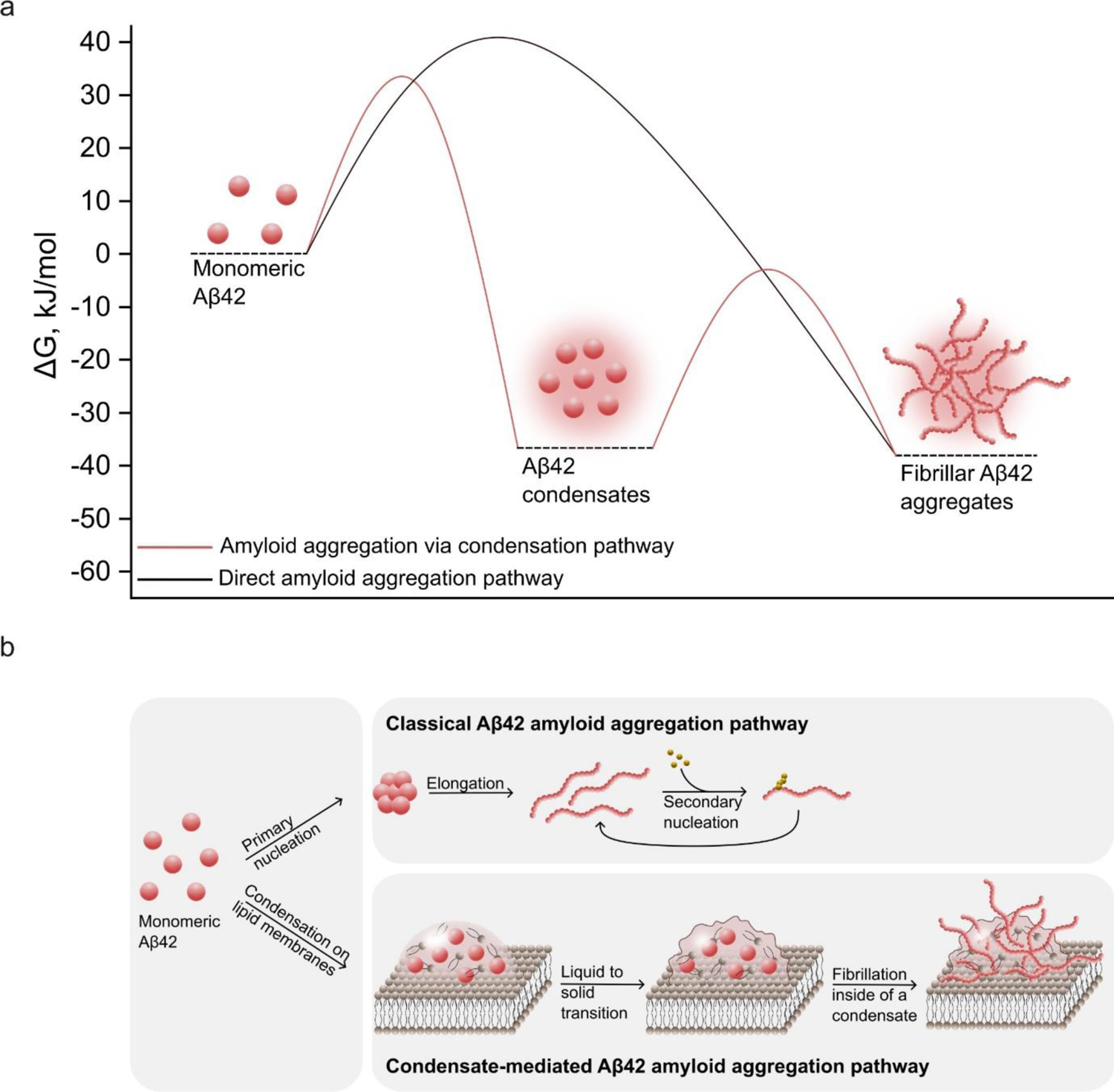
Aβ42 aggregates can form either through classical nucleation and growth or through condensation followed by a liquid-to-solid transition. a) Free energy diagram of monomeric, condensed and fibrillar Aβ42 states. b) Mechanisms of classical and condensate-mediated amyloid formation of Aβ42.

Crucially, we noted that the free energy barrier for the dilute monomer to dense liquid transition (ΔG^‡^_LLPS_ = 33.5 kJ/mol) surpasses that of monomer association into the condensates (ΔG^‡^(+) = 18 kJ/mol). This hints at an additional free energy barrier associated with condensate nucleation during the transition between the free monomeric and liquid phase separated states.

Overall, our investigation unveils that the energy barriers for Aβ42 LLPS on a lipid membrane are lower compared to direct Aβ42 fibril formation in solution. As such, the liquid-liquid phase separation followed by liquid to solid transition represents an alternative fibrillation pathway for Aβ42, which is energetically favoured in the presence of lipids through the emergence of the intermediate liquid phase (Figure 7b).

## Discussion

The crucial involvement of Aβ42 in AD has led to extensive exploration of its aggregation mechanisms both *in vivo* and *in vitro* (46–54). The late stages in the Aβ42 aggregation cascade are increasingly well understood. Once an aggregate nucleus is formed, it can grow further through elongation processes by which monomeric peptide molecules associate with the aggregated phases. The aggregates can also multiply either through fragmentation or through secondary nucleation (47,55). However, the mechanisms by which the original aggregate nuclei form have remained incompletely understood.

In our study, we used a set of fluorescence microscopy imaging methods, biophysical characterisation techniques and cell assays to uncover the existence of a biomolecular condensate phase of Aβ42 formed on lipid membranes both *in vitro* and in cellular systems. This liquid phase opens up a new, more energetically favourable aggregation pathway for Aβ42 by stabilizing the liquid state of Aβ42. Aβ42 co-phase separates with lipids and forms morphologically heterogeneous dynamic liquid condensates, which mature over time and ultimately form solid amyloid aggregates of high stability and low monomer dissociation rate. Our findings further suggest that Aβ42 condensates start hardening at the edge and gradually mature inwards to the liquid core. The Aβ42 liquid droplets are an environment with an increased local concentration of the peptide, and our results discussed in this paper show that the environment promotes the nucleation of the aggregated phase.

It is interesting to note that changes in lipid metabolism and lipid distribution in the brain that are commonly observed during the onset and development of AD (24,56–60), may directly affect Aβ42 phase behaviour, leading to altered amyloid plaque formation propensity and AD development. This idea is supported by the evidence of a high correlation between cholesterol and sphingolipid levels in the brain with AD progression and promoted Aβ42 aggregation and cytotoxicity by the latter (56–58). In addition, genetic mutations in proteins involved in brain lipid metabolism are the major risk factors for the development of sporadic forms of AD and the levels of omega-3 fatty acids are linked to a reduction of cognitive decline in AD and disease severity (61,62). Given that, a detailed understanding of the role of lipid chemistry in lipid-driven Aβ42 liquid-liquid phase separation mechanism is crucial in understanding the link between the changes in lipid membrane composition and AD progression.

In conclusion, our findings indicate that fundamental nucleation step in Aβ42 aggregation can be accelerated through the existence of a biomolecular condensate phase of the peptide. This type of two stage nucleation process may be a general pathway for neurodegenerative disease-related phase separating proteins, and evidence for this idea has been found for other systems such as FUS, TDP-43, α-synuclein, hnRNPA1 and tau (11,12,14,63–65). These ideas suggest the hypothesis that LLPS is a generic mechanism underpinning the amyloid disease pathology.

## Materials and Methods

### Protein expression, purification and labelling

Aβ42 WT and Aβ42 S8C mutant were expressed in *E. coli* and purified as described (66) except that 1 mM DTT was included in all buffers used for Aβ42 S8C purification. Before an experiment, lyophilised Aβ1-42 was dissolved in 6 M GuHCl 20 mM NaP 200 μM EDTA pH 8.5 buffer to solubilize any remaining aggregates, followed by a buffer exchange on a Superdex 75 Increase 16/30 column, in 20 mM NaP 200 μM EDTA pH 8.0 buffer. Eluted protein was collected from under the UV detector. The concentration of the unlabelled protein was determined using the (4) formula:

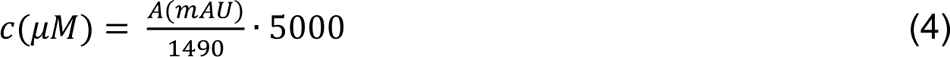

where A is the average recorded absorbance of the collected fractions at 280 nm with a 2 mm pathlength UV absorbance detector. The peptide was kept on ice throughout all stages.

Fluorescently labelled Aβ42 S8C mutant was prepared as described (67). Briefly, the peptide was solubilized in 6 M GuHCl 20 mM NaP 200 μM EDTA 1 mM DTT pH 8.0, buffer exchanged to 20 mM NaP 200 μM EDTA pH 8.0 and mixed with 4x molar excess of TAMRA maleimide dye. The protein-dye mix was incubated overnight at +4°C, followed by separation of the free dye the next morning. Free dye was separated by running two sequential size exclusions on a Superdex 75 Increase 16/30 column, in 20 mM NaP 200 μM EDTA pH 8.0.

### Formation of supported lipid bilayers

Chloroform solutions containing DMPC and cholesterol were mixed in glass vials. 1% β-BODIPY™ FL C12-HPC was added to label the liposomes. Chloroform was then evaporated with a gentle nitrogen stream, followed by the hydration of lipids with phosphate-buffered saline pH 7.4 (PBS). Lipid solutions were subjected to 5 freeze-thaw cycles in liquid nitrogen and a 42°C water bath. The vesicles were then extruded via a 50 nm pore size polycarbonate film track-etched membrane (Whatman, Cytiva) above the lipid phase transition temperature to obtain monodisperse liposomes, with a total of 31 passes. Supported lipid bilayers were then formed in PDMS observation chambers by depositing DMPC SUVs on cleaned glass slides (68).

### Confocal fluorescence imaging and fluorescence recovery after photobleaching

Confocal fluorescence imaging of Aβ42 on supported lipid bilayers (in PBS buffer) was performed in custom-made PDMS chambers containing supported lipid bilayers and bonded to a glass slide. The imaging was performed with a Leica Stellaris 5 confocal microscope equipped with a 63× oil immersion objective (Leica HC PL APO 63×/1.40 Oil CS2, NA 1.4). To perform fluorescence recovery after photobleaching experiments, an area of ca. 4 μm diameter was bleached with a laser line at 488 nm, followed by the recording of the images for 1–2 minutes every few seconds. The average fluorescence intensity of the area after photobleaching was then normalized to the average fluorescence intensity before and right after photobleaching. The fluorescence recovery half-times were calculated by fitting the data to the exponential growth model and using the formulas:

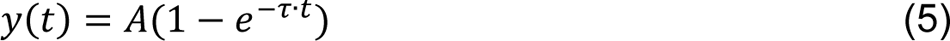

and

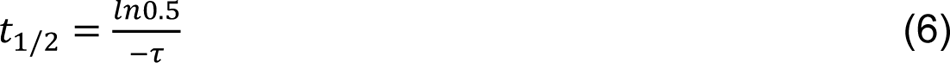

where y(t) is the average fluorescence intensity in the area of interest at time t, A is the plateau fluorescence intensity, τ is the fitted parameter and t_1/2_ is the recovery half-time.

### Total internal reflection microscopy imaging

Aggregation kinetics was monitored on a home-built TIRF microscope equipped with an autofocus system and a motorized XY stage as described previously (68). The samples were prepared in coverslip bottom four well chambers (Cellvis, USA). The chambers were cleaned using a 10 M aqueous solution of NaOH, followed by washing with 70% ethanol and then with MiliQ water. The plates were incubated first with the liposomes of DMPC (50 μM) for 15 minutes. Then 2 µM Aβ42 (final concentration) was added. The solutions were prepared in PBS buffer at pH 7.4 containing 5 mM β-mercaptoethanol (β-Me), 1 mM EDTA and 10 µM ThT. Aggregation was monitored by recording images at 25 fields of view in each sample chamber at a time interval of 30 minutes.

### SH-SY5Y culture and maintenance

Human neuroblastoma cells (SH-SY5Y) were cultured in DMEM/F-12 Glutamax™ (#10565018, Gibco) supplemented with 10% heat-inactivated fetal bovine serum (hiFBS, (#10082147, Gibco). Cells were kept at 37 degrees, 5% CO_2_ and 95% relative humidity.

### SH-SY5Y treatment with monomeric Aβ42

For hexanediol treatments, cells were plated in PerkinElmer Cell Carrier Ultra 96 well plates. For FRAP experiments, cells were plated in glass bottom slides (µ-Slide 8 Well Glass Bottom, Ibidi) coated with 0.002% w/v of polyornithine (PLO, #A-004, Sigma Aldrich) and 10 µg/mL of laminin (#L2020, Sigma Aldrich). After plating, cells were incubated overnight to allow adequate attachment.

30 minutes before the exogenous addition of monomeric Aβ42, the percentage of serum in the growth medium was decreased to 1%. Cells were then treated with freshly purified WT or TAMRA-labelled S8C Aβ42 at three different concentrations (0.5, 1 and 2 µM) and incubated for 0, 6, 16, 24 and 48h.

### Treatment with 1,6-hexanediol and cells fixation

At each time point, half of the growth medium was replaced with a freshly prepared 5% v/v 1,6-hexanediol solution (#240117, Sigma Aldrich) dissolved in a serum-free medium. Cells were incubated at RT for 2–3 minutes and washed once with warm DMEM/F-12. Cells were then fixed with 4% formaldehyde (#28906, Thermofisher) diluted in Dulbecco’s phosphate-buffered saline D-PBS (+/+) (#14040141, Gibco) and stored at 4°C.

### Immunocytochemistry

Cells were permeabilized with 0.1% Triton™ X-100 (#85111, Thermo Scientific) for 30 minutes at room temperature. Cells were then rinsed three times with D-PBS (+/+) and incubated for at least 1 h at RT in blocking solution (2% w/v of ultrapure BSA #9998, Cell Signalling, 3% v/v of goat serum in D-PBS (+/+)). No detergents were added to the blocking solution to avoid excessive permeabilization since that caused wheat germ-agglutinin dyes to stain intracellular compartments, rather than the outer cell membrane.

Afterwards, blocking solution was removed and cells were incubated with the APP/Aβ-specific primary antibody WO2 (#MABN10, Sigma Aldrich) at a 1:500 dilution for 16h at 4 °C. Cells were then washed three times with D-PBS (+/+) followed by incubation with Alexa Fluor™ 555 anti-mouse secondary antibody (#A-21422, Invitrogen) at a 1:1000 dilution for 1h at RT. All antibodies were prepared in a blocking solution unless otherwise stated. Cells were rinsed three times with D-PBS (+/+) and stained with Alexa Fluor™ 488 Phalloidin (#A12379, Invitrogen), Alexa Fluor™ 647 Wheat Germ Agglutinin (#W32466, Invitrogen) and Hoechst 33342 (#H3570, ThermoFisher). After 15 minutes of incubation at room temperature, cells were washed twice with D-PBS (+/+) and imaged using the Opera Phenix High-Content Confocal microscope. Image analysis and quantifications were performed with the Harmony High-Content Imaging and Analysis Software (Perkin Elmer).

### Calculation of the Gibbs free energy barrier of monomer association and dissociation to and from the condensates

In the transition from the initial state *x*_0_ to the final state *x*_*e*_, the transition rate constant k is described with the Arrhenius–Eyring equation (69):

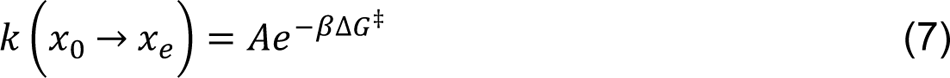

where ΔG^‡^ is the free energy barrier, β = 1/k_B_T and A is a pre-factor. We used the monomer diffusion-limited transition rate to approximate that the pre-factor A = D · r_eff_, where D = 1.4 ± 0.1 × 10^−10^ m^2^s^−1^ is the diffusion coefficient of free Aβ42 in solution. r_eff_ = 0.5 nm is the effective reaction volume of Aβ42 (45).

The relation between the kinetic rate constant k and ΔG^‡^ is described as follows (7):

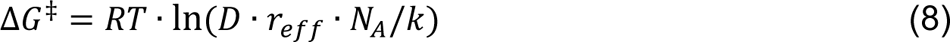

The rate constant k can be obtained from fitting the condensate growth kinetics data, assuming that the reaction order is 1:

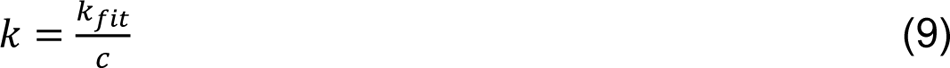

Where k_fit_ is the fitted monomer association rate parameter, obtained from fitting the condensate growth kinetics data with the formula:

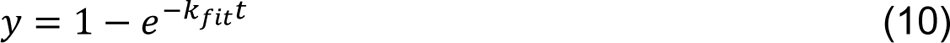

Where y is the associated monomer mass and t is time.

By combining equations no. 8 and 9, we express ΔG^‡^ for monomer association into condensates as:

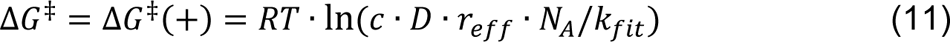

Free energy barrier associated with monomer dissociation from the condensates is calculated by adding the Gibbs free energy barrier for monomer association and the Gibbs free energy gain for monomeric to liquid transition:

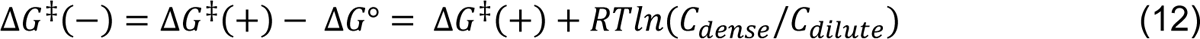

Where ΔG^‡^(+) is the free energy barrier for monomer association into condensates, R is universal gas constant, T is temperature and C_dense_/C_dilute_ is the ratio between concentration of free monomer in dense phase and dilute phase after the phase transition.

### Calculation of the Gibbs free energy barriers for protein phase transitions

The relation between the reaction time t and ΔG^‡^ is described as follows (7):

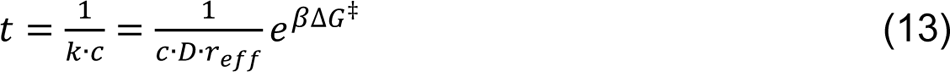

where β = (RT)^-1^, R is universal gas constant and T is temperature. c is the Aβ42 monomer concentration at the initial state. c = 2 μM in the direct monomeric to fibrillar transition and c = 40 mg/ml in the liquid-to-solid transition during LLPS. D = 1.4 ± 0.1 × 10^−10^ m^2^s^−1^ is the diffusion coefficient of free Aβ42 in solution. r_eff_ = 0.5 nm is the effective reaction volume of Aβ42 (45). Note that we estimated the prefactor A for the liquid-to-solid transition by using the parameters of free Aβ42 in solution. This choice corresponds to writing the rate constant using a phase-independent pre-factor and thereby partitioning the effect of interactions driving LLPS into the free energy barrier ΔG^‡^ (70). This enables us to meaningfully compare free energy barriers for amyloid formation, both with and without LLPS. Assuming the phase transition reaction order is 1, the Gibbs free energy barrier is then calculated from the reaction timescales:

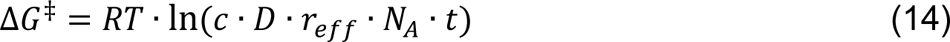

### Calculation of the Gibbs free energy gain during phase transitions

The Gibbs free energy gain was calculated as described in (41,42):

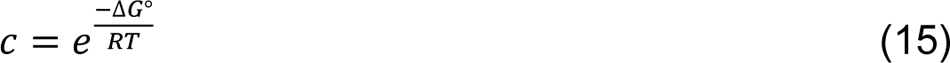

Which gives:

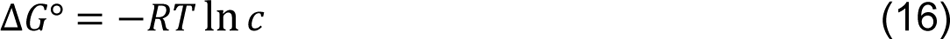

Where R is universal gas constant, T is temperature and c is concentration of free monomer in solution after the phase transition.

## Abbreviations

AD: Alzheimer’s disease

ALS: Amyotrophic lateral sclerosis

APP: Amyloid precursor protein

DMPC: 1,2-Dimyristoyl-sn-glycero-3-phosphocholine

FRAP: Fluorescence recovery after photobleaching

FUS: Fused in Sarcoma

LLPS: Liquid-liquid phase separation

PD: Parkinson’s disease

PDMS: Polydimethylsiloxane

TIRF: Total internal reflection fluorescence

## Acknowledgements

We thank Dr Aviad Levin for helpful discussions and advice. We also thank Becky Gregory, Ewa Andrzejewska and Lily Lin for help with Aβ42 S8C expression and purification. This work was funded by the UK Engineering and Physical Sciences Research Council (EPSRC) grant EP/S023046/1 for the Centre for Doctoral Training in Sensor Technologies for a Healthy and Sustainable Future (G.Š.), Fluidic Analytics Ltd (G.Š.). We would like to acknowledge funding from the European Research Council under the European Union’s Horizon 2020 research and innovation program through the ERC grant DiProPhys (agreement ID 101001615), the Frances and Augustus Newman Foundation (T.P.J.K.) and the Centre for Misfolding Diseases (G.Š., A.G.D., J.W., T.Š., M.V., T.P.J.K.). The Department of Atomic Energy, Government of India (project no. RTI 4007 to S.D.A and K.G.), the Science and Engineering Research Board, Government of India (grant no. CRG/2020/005527 to S.D.A and K.G.)

## Author contributions

G.Š., A.G.D., S.D.A., K.G., T.P.J.K. conceptualized and designed the study. G.Š., A.G.D., S.D.A. performed the experiments. T.Š. provided support with confocal imaging microscopy. G.Š., A.G.D., S.D.A., J.W., T.M. analyzed the data. All authors wrote and reviewed the manuscript.

