## Supplementary material for "The Alzheimer’s Aβ peptide forms biomolecular condensates that trigger amyloid aggregation": SI information

### Liquid-liquid phase separation on lipid membranes promotes Amyloid-β42 aggregation

This manuscript is available on biorXiv: https://www.biorxiv.org/content/10.1101/2024.01.14.575549v1

Classification: Biological Sciences – Biophysics and Computational Biology.

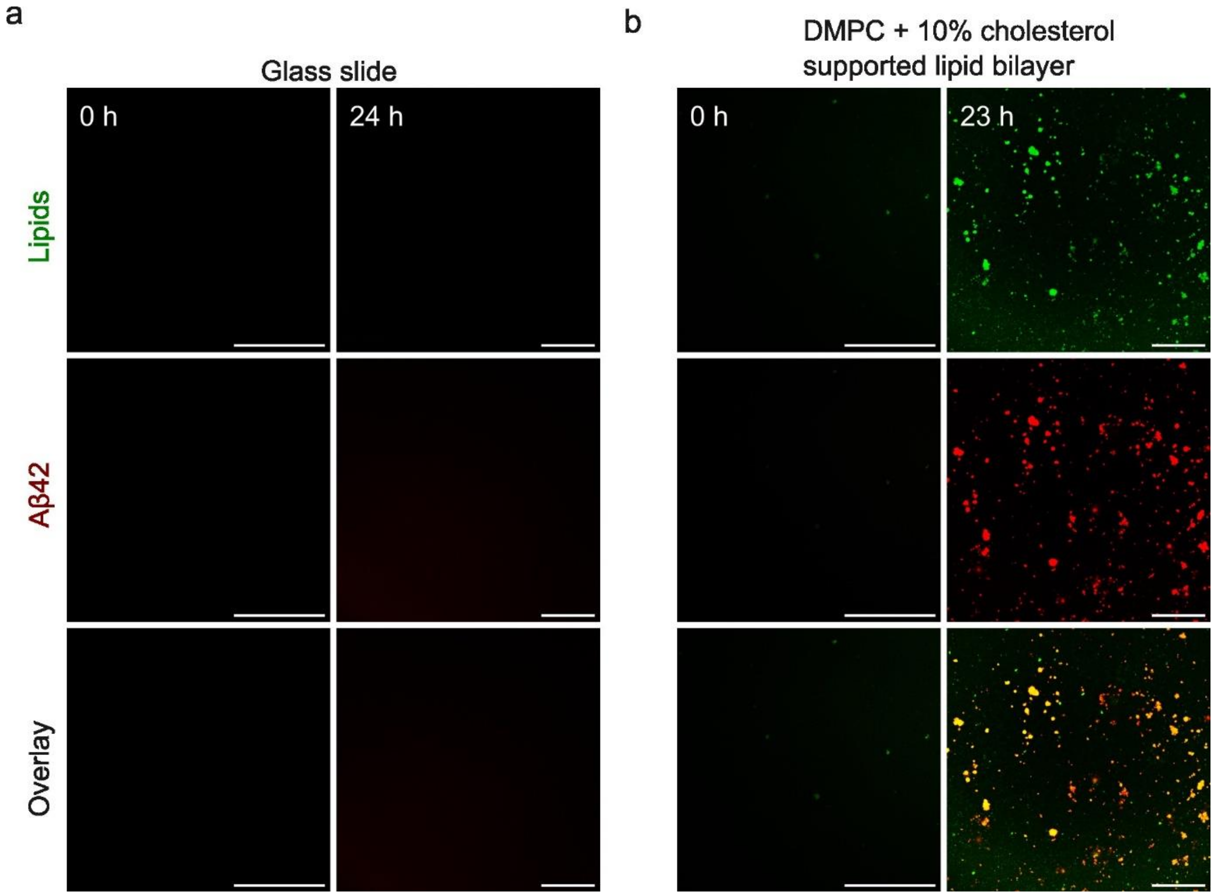


**Figure S1.** Aβ42 undergoes liquid-liquid phase separation on DMPC + 10% cholesterol supported lipid bilayers (b), but not on a glass surface (a). Scale bar = 50 μm.

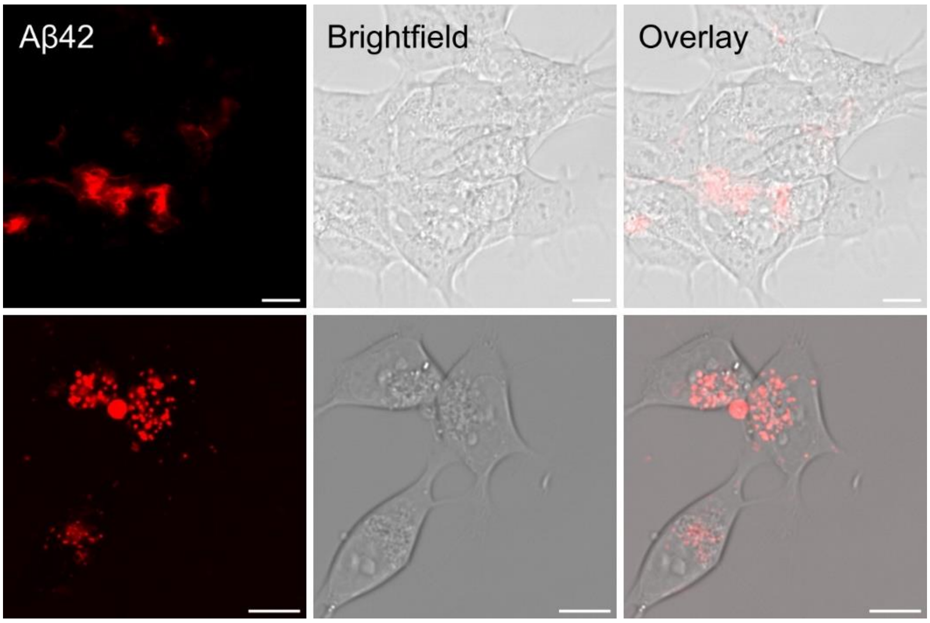


**Figure S2. Aβ42 aggregates colocalise with SH-SY5Y neuroblastoma cells.** Aβ42 staining: 5% Aβ42 S8C-TAMRA. Total protein concentration 2 μM. Scale bar = 10 μm. BF- bright field image.

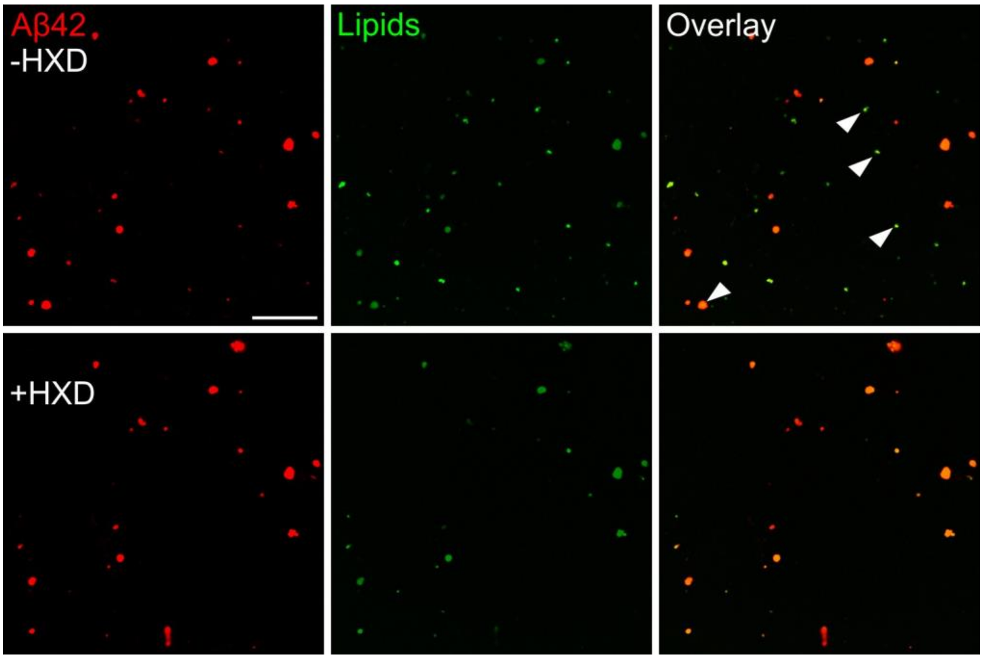


**Figure S3. Aβ42 and lipids inside of the condensates are resistant to 0.3 M (5%) 1,6-hexanediol treatment.** Confocal microscopy images Aβ42 condensates on DMPC + 10% cholesterol membrane, before and after treatment with 1,6-hexanediol. Samples were incubated for 21 h before the addition of 1,6-hexanediol. Scale bar = 50 μm.

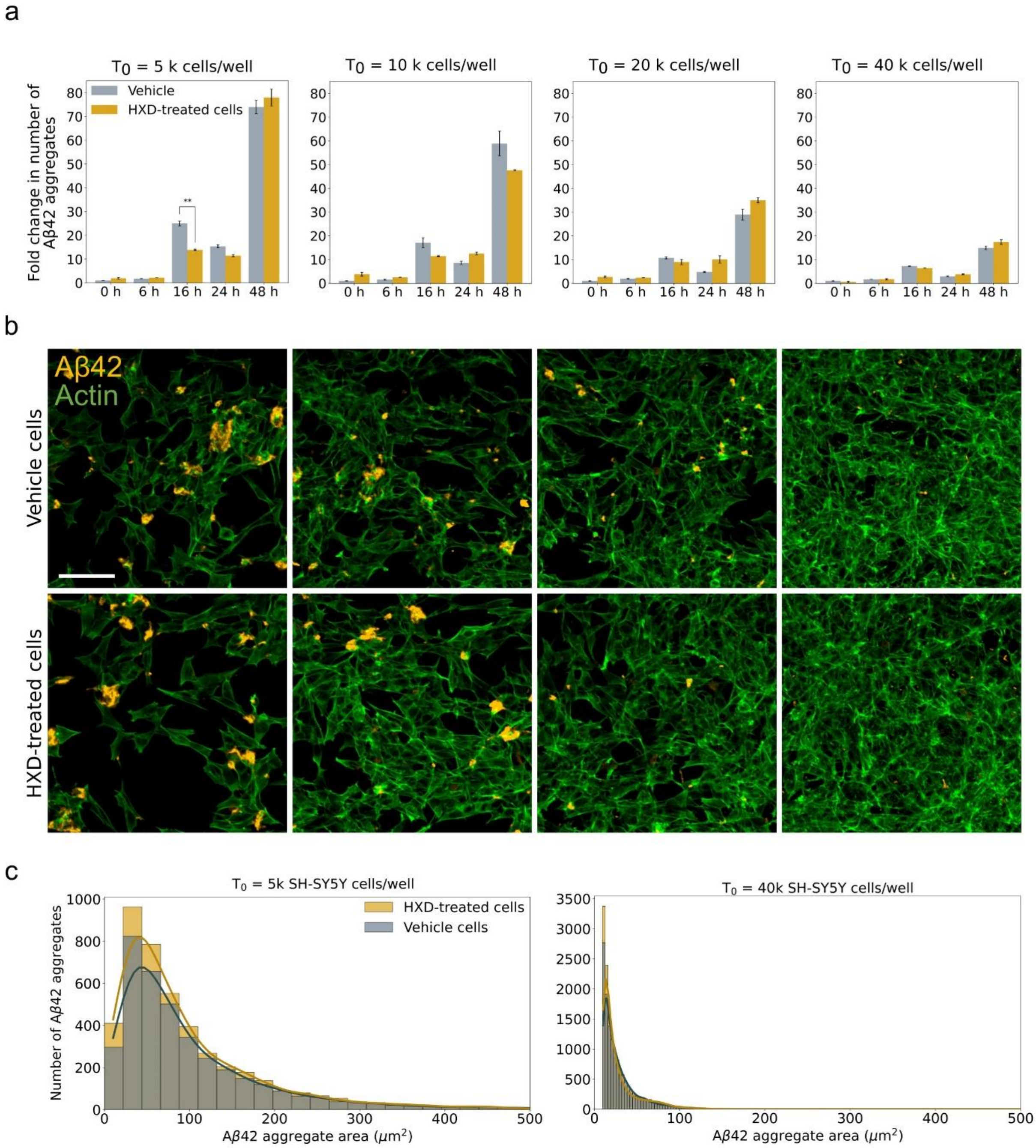


**Figure S4. Effect of cell density on Aβ42 condensate formation and aggregation.** a) fold change of the number of Aβ42 aggregates on SH-SY5Y cells over the number of Aβ42 aggregates on vehicle SH-SY5Y cells at t = 0 h. Cells were plated at four different densities at $T_0$ (5 000 (left), 10 000 (2nd from the left), 20 000 (3rd from the left) and 40 000 (right) per well) and fed with 2 μM of monomer. $T_0$ states for the day of plating (24 h before Aβ42 treatment). As cell density increases, both hexanediol-sensitive and resilient Aβ42 aggregate levels decrease. **indicate statistical significance with $p < 0.05$. b) Representative images of Aβ42 levels after 48 hours of monomer incubation on SH-SY5Y plated at four cell densities at $T_0$ (5 000 (left), 10 000 (2nd from the left), 20 000 (3rd from the left) and 40 000 (right) per well). Scale bar = 100 μm. c) Size distribution of Aβ42 aggregates at low (5 000/well) and high (40 000/well) cell densities after 48 hours of monomer incubation. Larger aggregates are observed at low cell densities (left panel), whereas smaller aggregates are observed at high cell densities (right panel).

Aβ42 species are resilient to 1,6-hexanediol independently of the cell density, indicating maturity of the aggregates at the later time points. Lines show kernel density estimate plots.

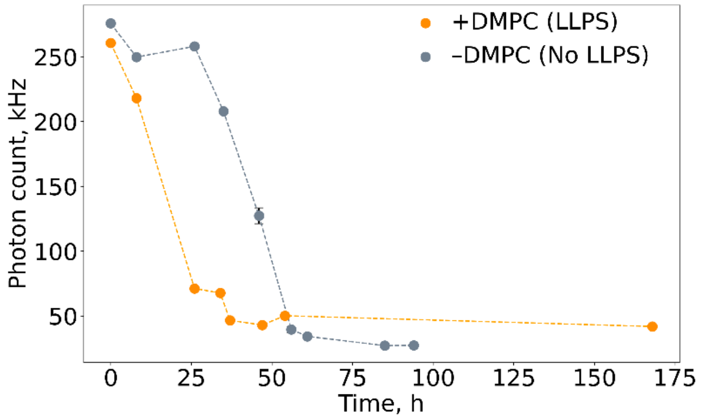


**Figure S5.** FCS photon count of soluble Aβ42 under LLPS conditions (on DMPC supported lipid bilayer) and under direct fibrillation conditions (on a glass slide) over time. Dashed lines are guides to the eye.

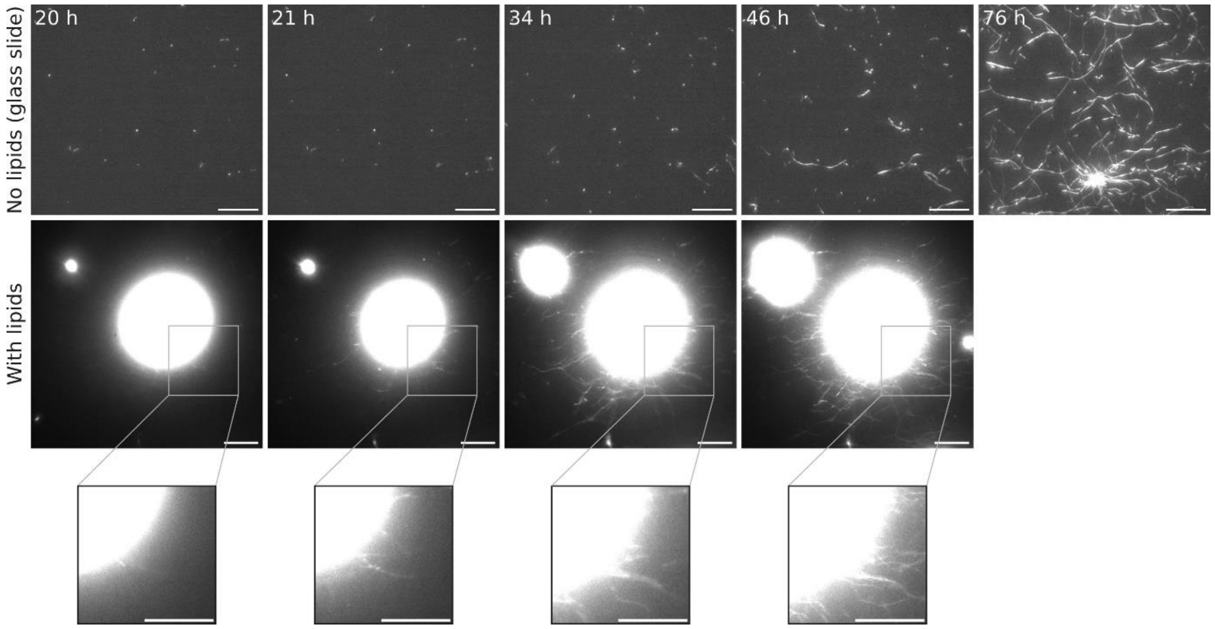


**Figure S6**. Condensation triggers  Aβ42 fibril formation. TIRF microscopy images of Aβ42 fibrils forming on a glass slide (non-LLPS conditions) and on a DMPC supported lipid bilayer (LLPS conditions). Under non-LLPS conditions, (in the absence of lipids), fibrils do not form until 46 hours (top row). Under LLPS conditions, condensate-associated fibrils form after 21 hours of incubation. 2 μM Aβ42 was used in the experiments. Aggregates were stained with ThT. Scale bar = 10 μm.

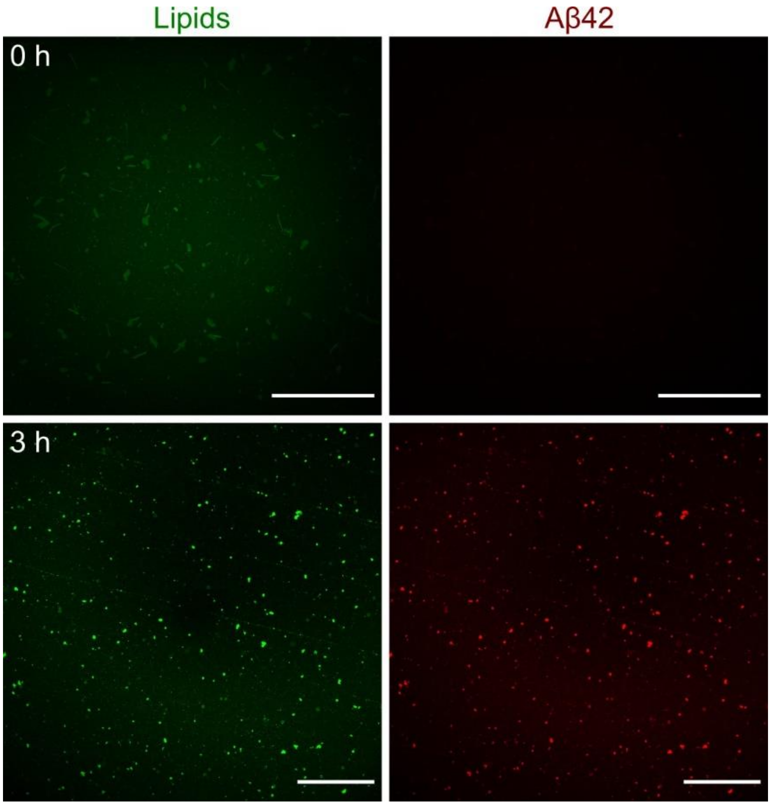


**Figure S7.** Confocal fluorescence microscopy images of Aβ42 condensates on DMPC + 10% cholesterol lipid membrane at t = 0 h and t = 3 h. Scale bar = 50 µm.

### Supplementary Methods

#### Estimation of diffusion coefficient of free Aβ42 in solution

Diffusion coefficient of Aβ42 in solution was calculated using the Stokes-Einstein equation (1):

$$D = \frac{k_B T}{6\pi \eta r} \tag{1}$$

Where $k_B$ – Boltzmann constant (J/K), T – temperature (K), ŋ – dynamic viscosity (Pa·s), r – hydrodynamic radius of a particle (m). The hydrodynamic radius of Aβ42 is measured by MDS as described in (1).

#### Estimation of diffusion coefficient of Aβ42/lipids in a condensate

Diffusion coefficient (D) of Aβ42/lipids inside of a condensate was estimated from FRAP data using a Soumpasis equation (2) (2):

$$D = 0.224 \frac{r^2}{t_{1/2}} \tag{2}$$

Where r – radius of the bleached area, $t_{1/2}$ – FRAP recovery half-time.

**Fluorescence correlation spectroscopy**

The home-built TIRF setup was modified to enable FCS measurements in the second emission port of the microscope. A multimode optical fiber of core diameter of 50 µm serves as the confocal pinhole, and also carries the photons to the avalanche photodiode single photon counting detector (MPD, Italy). A solid state 488 nm laser was used for fluorescence excitation of the ATTO 488-labelled Aβ42 or the BODIPY-PC. A digital correlator card (Becker and Hickl, Germany) was used for calculation of the autocorrelation functions. The condensates were first visualized by imaging, then the sample stage was moved appropriately to position the point of interest at the focus for FCS measurements. The FCS autocorrelation data were fitted using a one-component diffusion model as described in (4). The concentration of free Aβ42 protein in the soluble fraction was calculated from the analysis of the FCS autocorrelation curves and the intensity time traces of the ATTO 488-labelled Aβ42.
